# Vegetative compatibility groups explain variation in the virulence of *Verticillium dahliae* on strawberry

**DOI:** 10.1101/181172

**Authors:** Rong Fan, Helen Cockerton, Andrew D. Armitage, Helen Bates, Emma Cascant-Lopez, Laima Antanaviciute, Xiangming Xu, Xiaoping Hu, Richard J. Harrison

## Abstract

*Verticillium dahliae* infection of strawberry (*Fragaria × ananassa*) is a major cause of disease-induced wilting in soil-grown strawberries across the world. To understand what components of the pathogen are affecting disease expression, the presence of the known effector *VdAve1* was screened in a sample of *Verticillium dahliae* isolates. Isolates from strawberry were found to contain *VdAve1* and were divided into two major clades, based upon their vegetative compatibility groups (VCG); no UK strawberry isolates contained *VdAve1*. VC clade was strongly related to their virulence levels. *VdAve1*-containing isolates pathogenic on strawberry were found in both clades, in contrast to some recently published findings.

On strawberry, *VdAve1*-containing isolates had significantly higher virulence during early infection, which diminished in significance as the infection progressed. Transformation of a virulent non-*VdAve1* containing isolate, with *VdAve1* was found neither to increase nor decrease virulence when inoculated on a susceptible strawberry cultivar. There are therefore virulence factors that are epistatic to *VdAve1* and potentially multiple independent routes to high virulence on strawberry in *V. dahliae* lineages.

Genome sequencing a subset of isolates across the two VCGs revealed that isolates were differentiated at the whole genome level and contained multiple changes in putative effector content, indicating that different clonal VCGs may have evolved different strategies for infecting strawberry, leading to different virulence levels in pathogenicity tests. It is therefore important to consider both clonal lineage and effector complement as the adaptive potential of each lineage will differ, even if they contain the same race determining effector.

## Introduction

### *V. dahliae* hosts and control methods

*Verticillium dahliae* Kleb. is a soilborne plant pathogenic fungus which can cause wilting on over 200 plant species, including many agricultural and horticultural crops [1]. This pathogen has a rapidly expanding host range, most notably on the crops lettuce and cauliflower where it has become a major pathogen within these industries [2]; [3,4]. *V. dahliae* can not act as an effective saprophyte and thus must produce resting structures to persist within the soil between infection events. The resting structures are microsclerotia, these are robust melanised hyphae which are very difficult to eradicate and can persist in the soil for up to 20 years post-formation [1]. Soil fumigation is the only effective control measure however it relies on the use of chemicals with harmful environmental effects which have either been banned or (in the EU/UK) are due to be phased out [5]. Furthermore, crop rotation is largely ineffective as a management strategy and there are no available fungicides to combat the disease once established [6,7]. Consequently, breeding natural host disease resistance into crops remains the most effective strategy to control Verticillium wilt disease.

### The Ve1 resistance gene is operational against ‘race 1’ isolates

Pathogens have evolved to deliver effector proteins in order to suppress host PTI (pathogen associated molecular pattern [PAMP]-triggered immunity). If successful, the pathogen will cause effector-triggered susceptibility (ETS) in the host and as such allow the pathogen to cause disease [8]. The plant cell wall is the primary barrier preventing pathogen infection. In order to overcome this barrier, pathogens secrete cell wall-degrading enzymes (CDEs) which are able to catabolise plant cell wall polysaccharides [9]. Although CWDEs are pathogenicity genes [9–11], some also function as PAMPs, and trigger host PTI independent of their depolymerisation activity [9–11].

Plants have an innate immune system, part of which is comprised of so called ‘resistance’ genes that encode receptors to recognize molecular patterns of pathogen proteins [12] and effectors [13]. Dominant plant resistance genes that bring about resistance to *V. dahliae* ‘race 1’ isolates have been identified in cotton, lettuce, tomato, and sunflower [14–17] have been deployed successful in commercial tomato (Ve1) [16], and lettuce (Vr1) [18] cultivars. Ve1 is a cell surface like receptor operative against ‘race 1’ of the pathogen (defined by resistance in tomato), which carries the avirulence gene *VdAve1* [19], [20]. However, Ve1 on tomato and its orthologs in other crops are not effective against ‘race 2’ isolates which lack *VdAve1* [21]. Although attempts have been made to find tomato cultivars resistant to race 2 isolates, no high level resistance has been discovered [22].

### Race structure in current crops

It was thought that genome rearrangement was responsible for the widespread success of *V. dahliae* as a pathogen across multiple host species [23]; [24], however, a similar pattern of rearrangement has been observed across the 10 species within the Verticillium genus, including those which are plant pathogens with a limited host range [25]. It is now proposed that gene loss is more important in defining differences in speciation and phenotypic diversity within the *Verticillium spp.* [25].

*V. dahliae* has linage specific genome regions enriched in effectors which have arisen by segmental duplication to allow for functional divergence [24–26]. By contrast, the effector *VdAve1* was most likely acquired by horizontal gene transfer from the host plant on a single occasion and this effector has undergone multiple independent gene loss events within different ‘race 2’ isolates of *V. dahliae* [20]; [27]. Indeed, *VdAve1* is flanked by several ‘young’ transposable elements in a lineage specific region of *V. dahliae*, which, it is believed provide a mechanism of gene loss [20]; [27,28]. Furthermore, it was suggested that ‘race 2’ isolates should be divided into discrete groups based on the sequences in the genomic region around *VdAve1* gene [24] with these regions representing each *VdAve1* loss event. Greater complication of the race structure has arisen, whereby ‘race 2’ isolates infecting tomato in Japan can be split into two further races based on their differential pathogenicity on resistant rootstocks [29], however further work is required to determine the causative effector(s).

A study investigating race distribution in Californian tomato exclusively isolated ‘race 2’ strains from tomato, despite isolating ‘race 1’ strains from other crop hosts. It was proposed that this is due to the predominant use of Ve1 containing tomato cultivars [30]. Likewise, *V. dahliae* strains from cotton, mint, olive, and potato were exclusively found to be ‘race 2’ isolates whereas all isolates from artichoke were ‘race 1’ and isolates from lettuce, pepper and Californian strawberry contained both races [31]. It is clear that there is a global overlapping distribution on ‘race 1’ and ‘race 2’ isolates however, the host determines the proportion of the races isolated [31].

A complex race structure in strawberry has been defined through a series of pathology experiments by Govorova and Govorov (1992), who identified six newly described races of Russian *V. dahliae* isolates based on resistance associated with five cultivars [32]. This work did not seek to quantify race structure in terms of *VdAve1* presence nor the effector complexes however there is support for a complex series of plant- pathogen interactions. Strawberry wilt resistance was identified at the QTL level, in wilt segregating progenies planted within a mixed inoculum field, resistance was not monogenic, but quantitative, thus providing further evidence for complex plant- pathogen interaction [33].

*V. dahliae* is an asexual organism comprised of several discrete clonal lineages. Vegetative compatibility groups (VCGs) determine whether *V. dahliae* strains can form hyphal anastomosis bridges, leading to genetic exchange through nuclear fusion and subsequent haploidisation. VCGs limit the transmission of genetic material between unrelated strains and thus promote the maintenance of distinct clonal lineages [34]. Previous studies have described two VCGs present on UK strawberry: VCG 2A and VCG 4B [35]. However recent work has redefined the clonal lineages based on over 26,000 SNPs and as such divided the population structure of *V. dahliae* into two major clades (I & II) with clade II dissected into two further sub clades (II-1 & II-2) [36].

### (4) Objective of this study

The objective of this study was to characterise whether UK strawberry cultivars were infected by both ‘race 1’ and ‘race 2’ isolates of *V. dahliae* and if so whether *VdAve1* played any role in virulence or avirulence.

## Materials and Methods

### PCR detection of VdAve1 gene in V. dahliae isolates

DNA of *V. dahliae* isolates was extracted using Macherey–Nagel Plant DNA Extraction Kit (Germany) following the manufacturer’s instruction. Primers targeting a partial region of the *VdAve1* open reading frame, VdAve1-F/R (Supplementary Table 1), were designed using Geneious 7.0 and the *V. dahliae* specific primers Vd-F929-947/Vd-R1076-1094 (Supplementary Table 1) were used as a control to confirm DNA quality (Wei et al. 2015). The DNA of a *VdAve1*-carrying isolate (*V. dahliae* 12067) was used as a positive control. PCR conditions were 95 °C for 5 min, followed by 35 amplification cycles of 30 s at 95 °C, 30 s at 60 °C and 30 s at 72 °C. Amplicons were separated on a 2 % agarose gel.

**Table 1.** Strains used in this study

| Isolate | Species | Host | Origin | Date isolated | Ave1 | VCG |
| --- | --- | --- | --- | --- | --- | --- |
| 12008 | *V.dahliae* | Strawberry | Kent, UK | 1985 |  | 2B |
| 12009 | *V.dahliae* | Strawberry | UK | 1986 |  |  |
| 12023 | *V.dahliae* | Strawberry | Warwickshire, UK | 1990 |  |  |
| 12024 | *V.dahliae* | Strawberry | Norfolk, UK | 1990 |  | 2A |
| 12025 | *V.dahliae* | Strawberry | Norfolk, UK | 1990 |  |  |
| 12027 | *V.dahliae* | Strawberry | Kent, UK | 1990 |  |  |
| 12029 | *V.dahliae* | Strawberry | Norfolk, UK | 1990 |  | 2A |
| 12030 | *V.dahliae* | Strawberry | Hereford, UK | 1990 |  | 2B |
| 12031 | *V.dahliae* | Strawberry | Somerset, UK | 1990 |  | 2A |
|  |  |  | West Midlands, |  |  |  |
| 12033 | *V.dahliae* | Strawberry | UK | 1990 |  |  |
| 12045 | *V.dahliae* | Acer | Kent, UK | 1993 |  | 2A |
| 12046 | *V.dahliae* | Acer | Kent, UK | 1993 | + | 2A |
|  |  |  | Worcestershire, |  |  |  |
| 12047 | *V.dahliae* | Cotinus | UK | 1992 | + | 2B |
|  |  |  | Worcestershire, |  |  |  |
| 12048 | *V.dahliae* | Cotinus | UK | 1992 |  |  |
| 12049 | *V.dahliae* | Acer | Kent, UK | 1993 |  | 2A |
| 12050 | *V.dahliae* | Acer | Beds, UK | 1993 | + | 2B |
| 12051 | *V.dahliae* | Hop | Kent, UK | 1987 | + | 2B |
| 12052 | *V.dahliae* | Acer | Suffolk, UK | 1992 | + | 2B |
| 12056 | *V.dahliae* | Strawberry | Kent, UK | 1985 |  | 2A |
| 12057 | *V.dahliae* | Strawberry | Kent, UK | 1985 |  |  |
| 12059 | *V.dahliae* | Phlox | Kent, UK | 1986 | + |  |
| 12064 | *V.dahliae* | Chrysanthemum | Bristol, UK | 1968 |  |  |
| 12067 | *V.dahliae* | Tomato | Dorset, UK | 1971 |  |  |
| 12085 | *V.dahliae* | Raspberry | Kent, UK | 1998 |  | 2B |
| 12086 | *V.dahliae* | Strawberry | Kent, UK | 1988 |  |  |
| 12087 | *V.dahliae* | Strawberry | Kent, UK | 1988 |  | 2B |
| 12088 | *V.dahliae* | Strawberry | Unknown | 1998 |  | 2B |
| 12089 | *V.dahliae* | Acer | Kent, UK | 1998 | + | 2B |
| 12099 | *V.dahliae* | Hop | Kent, UK | 1998 |  | 2A |
| 12100 | *V.dahliae* | Hop | Kent, UK | 1998 | + | 2A |
| 12125 | *V.dahliae* | Strawberry | Kent, UK | 1989 |  |  |
| 12152 | *V.dahliae* | Strawberry | Belgium | 1996 |  |  |
| 12158 | *V.dahliae* | Strawberry | Kent, UK | 2000 |  | 2A |
| 12161 | *V.dahliae* | Strawberry | Lincolnshire, UK | 2000 |  | 2A |
| 12163 | *V.dahliae* | Strawberry | Norfolk, UK | 2000 |  | 2B |
| 12223 | *V.dahliae* | Hop | UK | 1998 | + | 2A |
| 12251 | *V. dahliae* | Strawberry | Kent, UK | 2012 |  | 2B |
| 12253 | *V.dahliae* | Strawberry | Kent, UK | 2012 |  | 2B |
| USA- |  |  |  |  |  |  |
| 07001 | *V.dahliae* | Strawberry | USA | NA |  | 2A |
| USA-15- |  |  |  |  |  |  |
| 081 | *V.dahliae* | Strawberry | USA | NA |  | 2A |
| USA-15- |  |  |  |  |  |  |
| 082 | *V.dahliae* | Strawberry | USA | NA |  | 2A |
| USA-V1 | *V.dahliae* | Strawberry | USA | NA |  | 2A |

### PCR detection of vegetative compatibility group

The vegetative compatibility group of the isolates used within the pathogenicity assay was determined through PCR amplification of a lineage specific region using the primers DB19 and DB22 [35,37]. The PCR was conducted using Q5 High-Fidelity DNA polymerase with conditions of 98 °C 30 s; 35 cycles of 98 °C 10 s, 55 °C 30 s, 72 °C 30 s and 72 °C at 2 min. Amplicon production was confirmed through gel electrophoresis on a 1 % agarose gel before PCR clean up using the Machery-Nagel Nucleospin® Gel and PCR clean up kit and sequenced in two directions using Sanger sequencing.

### Phylogeny of VC groups

Multiple sequence alignment of the region amplified by the DB19/22 primer set was performed using MUSCLE [38]. A neighbour-joining phylogeny was constructed using the HKY85 substitution model, with support from 1000 Bootstrap replicates in the Geneious v10 software package (http://www.geneious.com) [39]. The resulting dendrogram was visualised using FigTree (http://tree.bio.ed.ac.uk/software/figtree/).

### Pathogenicity assay on strawberry

The wilt susceptible strawberry cv. Hapil was used to determine the pathogenicity of isolates. Conidial inoculum of thirteen isolates were prepared through transfer of five, 1 cm plugs of mycelia growing on PDA into 50 ml of PDB media incubated in the dark at 25 °C and 150 rpm for one week. After filtration through 13 μm particle retention filter paper, the conidial density was adjusted to 1x10^6^ spores.ml^−1^ using sterile distilled water. Three-week strawberry runners were up-rooted from pots, excess soil was removed with water, old and trimmed 2 cm from the root tips. Roots were dipped in spore suspensions for 5 min and replanted in the pot. Plants were kept in a growth room with 16:8 hr light - dark cycles and at a constant temperature of 22 °C. Disease severity was surveyed following five categories: 0 healthy shoots, all leaves green; 1 - single leaf yellow-brown; 2 - <25% leaves yellow-brown; 3-26–50% leaves yellow-brown; 4-51–75% leaves yellow-brown; 5-76–100% leaves yellow-brown, the plants dead [40].

### Statistical analysis

Wilt severity scores (*Y*_*i*_) [= 0, 1, 2, 3, 4 and 5] were analysed using a cumulative logit model in which *Y*_*i*_ is assumed to follow a multinomial distribution. A cumulative logit model is a regression model for a cumulative logit [41]:

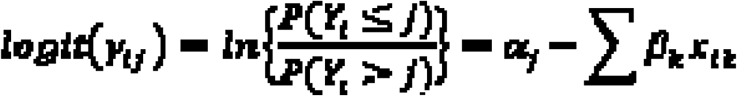

where *x*_*k*_ represents the k^th^ explanatory variate, the *ß*_*k*_ effect of *x*_*k*_, and α_*j*_ the intercept for each cumulative logit. The larger the value of 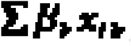, the higher the probability of *Y*_*i*_ falling in a category at the upper end of the category scale. It is the extension of the common logistic model with only two possible outcomes (i.e. diseased or healthy). *ß* estimates the change in the cumulative odds ratio (on log_e_ scale) for one unit increase in the explanatory variate *x*. In the present study, there were three categorical explanatory variates: VCG, host and race group.

### Generation of VdAve1 knock-in mutants and induction of VdAve1 expression

The known avirulence gene gene *VdAve1* and its native promoter *(called PAve1)* were cloned from the ‘race 1’ isolate 12067 using the primer pair VdAve1C-F/R and NPAve1-F/R, respectively (Supplementary Table 1). The *VdAve1* random insertion vector was constructed using USER-Brick vector system, [42]. Core USER Bricks, as well as the constitutive fungal promoter *PgpdA* and the selective marker *Hygromycin,* were amplified from the plasmid pRF-HU2-F/R and pRF-HUE-F/R using the primers specified in [42]). Two different insertion vectors were assembled; one containing the *Ave1* native promoter, and another using the constitutive promoter *PgpdA.* Correct vector assembly was then verified by PCR. The valid vectors were transformed into *Agrobacterium tumefaciens* isolate GV3101. After *A. tumefaciens*-mediated transformation (ATMT) of *V. dahliae*, the hygromycin B-resistant transformants were purified through the single-spore purification process as described previously [43]. Amplicon insertion in the *VdAve1* knock-in isolates was confirmed through PCR with the primer pairs Hyg-F/R and VdAve1-F/R (Supplementary Table 1) and qRT-PCR detection with the housekeeping genes of elongation factor 1-α (EF-F/R) and ß-tubulin (btubulin-F/R), as well as the specific primers for *VdAve1* (qAve1-F/R, Supplementary Table 1). Moreover, in order to induce *VdAve1* expression, cultures of five *VdAve1* knock-in mutants of each vector were incubated on PDA and Capek-Dox agar media for 5 days at 25°C before qRT-PCR quantification.

### Genome assembly and annotation

Genome sequencing was performed on five *V. dahliae* isolates. For isolates, 12251, 12153, 12158 and 12161 DNA extraction was performed on freeze-dried mycelium using a GenElute plant DNA miniprep kit (Sigma-Aldrich). Paired-end genomic libraries were prepared using a Nextera sample preparation kit (Illumina), according to the manufacturer’s protocol. Analysis using a fragment analyzer (Advanced Analytical Technologies) confirmed that libraries had a high representation of DNA fragments 600-1,000 bp in length. Paired end libraries were sequenced using 250 bp reads on an Illumina MiSeq machine. Genomic DNA for isolate 12008 was prepared by extracting high molecular weight DNA from freeze-dried V. dahliae mycelium using the Macherey-Nagel Nucleospin Plant II kit (Fisher 11912262) for Illumina sequencing and QIAGEN Genomic-tip 500/G kit(Qiagen,Germany) for Pacbio sequencing, which was sequenced on an RSII machine. An Illumina library was also prepared for isolate 12008, with DNA sheared using the Covaris M220 with microTUBE-50 (Covaris 520166) and size selected using a Blue Pippin (Sage Science). The Illumina library was constructed with a PCR-free method using NEBNext End Repair (E6050S), NEBNext dA-tailing (E6053S) and Blunt T/A ligase (M0367S) New England Biolabs modules. The library was sequenced using a 2x 300 bp PE (MS-102-3003) kit on an Illumina Miseq v3.

Illumina data adaptor sequences and low-quality data were removed using fastqc-mcf (available from https://expressionanalysis.github.io/ea-utils/) and the output was assembled with SPAdes [44]. RepeatMasker and TransposonPSI were used to identify repetitive sequence regions [45] [46]. Long reads generated from Pacbio sequencing of isolate 12008 were assembled using Canu and polished using Illumina MiSeq reads in Pilon to correct erroneous SNPs and InDels [47,48]. The assembly was edited in accordance with recommendations from the NCBI contamination screen (run as part of submission to Genbank in November 2016 for isolate 12008 and July 2017 for illumina-only isolates) with contigs split, trimmed or excluded as required. Quast [49] was used to summarise assembly statistics and BUSCO [50] was used to assess completeness of gene space within the assembly. Gene prediction was performed using Braker1 and CodingQuarry [51,52] and functional annotations predicted for these proteins using InterproScan [53]. Sequence data and annotated genomes were deposited at DDBJ/EMBL/GenBank within Bioprojects PRJNA344737 (isolate 12008) and PRJNA352681 (isolates 12151, 12153, 12158 and 12161).

For newly sequenced genomes and for the existing JR2, VdLs17 [54] and Va.Ms.102 [28], putative virulence factors and effector candidates were identified within predicted gene models. Secreted proteins were identified using SignalP 4.1 [55], removing those that were predicted to contain a transmembrane domain (TMHMM) or a GPI anchor (GPIsom). Small secreted cysteine-rich proteins (SSCP) were identified from secretomes, detecting proteins shorter than 300 amino acid and having a cysteine content of greater than 3 %. Secreted cell wall degrading enzymes were identified within from those proteins predicted as carbohydrate active enzymes (CAZYmes) from the CAZY database [56]. Further effector candidates were identified in the predicted secretome using EffectorP [57]. LysM proteins were identified using CAZY hmm model CBM50.hmm. NPP1 proteins were identified from Interproscan function annotations (IPR008701).

### Generation of a genome-wide phylogenetic tree

In order to study the genomic region surrounding the *VdAve1* locus, Illumina MiSeq reads of the five UK ‘race 2’ isolates were mapped to the reference genome of JR2 using Bowtie2 and the alignment visually inspected using IGV [58]. The phylogenetic tree of these five ‘race 2’ isolates along with the isolates of VdLs17 and JR2 was generated using RealPhy [59] using *V. dahliae* isolate JR2 as a reference genome.

### Orthogroup analysis

Gene expansion was investigated through clustering proteins into ortholog groups and then identifying the number of genes from each organism within that orthogroup. The proteomes of the five sequenced isolates and reference isolates JR2, VdLs17 and VaM102 were clustered using OrthoMCL [60] using a cut-off length of 50 amino acids and an inflation value of 5.

## Results

### UK strawberry isolates do not contain *VdAve1*

In order to investigate whether there is any evidence for the presence of *VdAve1* in *V. dahliae* in the UK, we screened a total of 38 isolates isolated from UK strawberry, hop, acer, raspberry, potato, phlox, cotinus, tomato and chrysanthemum by PCR using a pair of primers targeting a 351 bp region within the *VdAve1* gene (see Table 1 and Supplementary Table 1). In addition primers of Vd-F929-947/ Vd-R1076-1094 targeting part of *V. dahliae* intergenic spacer regions, were used as a positive control for DNA quality and PCR success in all DNA samples, generating a band of 160 bp. In addition, four *V. dahliae* isolates from Californian strawberry population were also included to confirm the previously reported presence of *VdAve1* [31]. In the UK isolates, the *VdAve1* gene was present in hop, acer and cotinus isolates indicating the presence of ‘race 1’ *V. dahliae* within the UK (Supplementary Fig. 1A). No DNA amplification at the *VdAve1* locus was observed in the 23 *V. dahliae* isolates from UK strawberry, (Supplementary Fig. 1B). We found that the Californian strawberries isolates contained both races, confirming previous studies [31]. These results show that ‘race 1’ strawberry isolates do exist but are absent among isolates from UK strawberry, despite the presence of *VdAve1*-containing isolates from other hosts within the UK. The observed frequency of *VdAve1* containing UK isolates across other hosts is 9/16 (Supplementary Fig. 1A).

**Fig 1.**
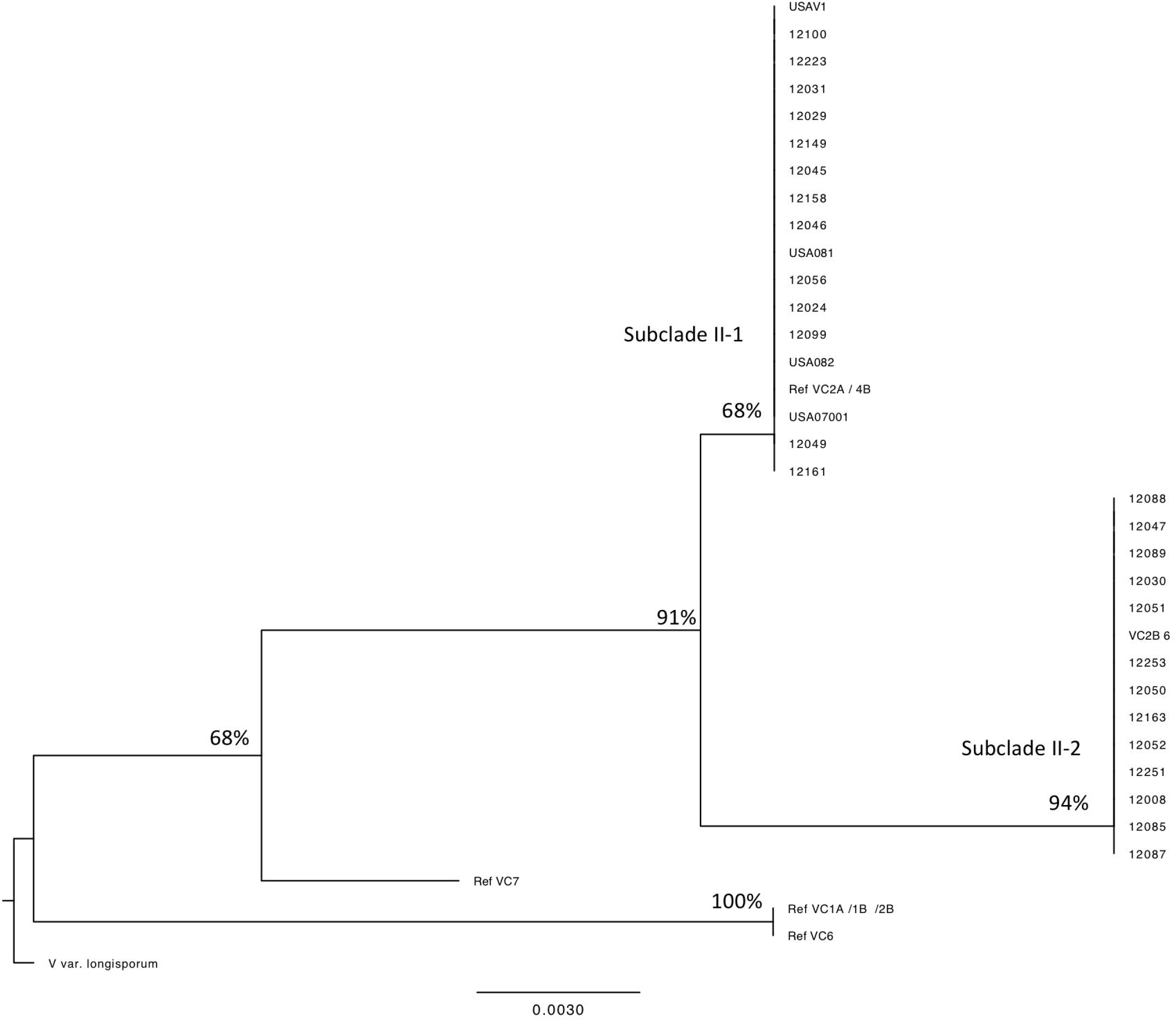
Neighbour joining phylogeny of *Verticillium dahliae* using primers DB19/22 to classify VC clades. Reference sequences were obtained from Collado-Romero et. al [35]. Bootstrap support values are the result of 1000 resampling events.

### *VdAve1* is not associated with a single VC group

Further characterisation of a subset of 30 isolates from the the initial screen for *VdAve1* revealed variation in vegetative compatibility groups of the isolates in this study. Contrary to recent, but not all reports, *VdAve1* was found in two major VC clades- VC subclade II-1 and VC subclade II-2 (Fig. 1) and not just a single VCG. The two VC clades identified in this study contain the previously reported VC groups on UK strawberry: VGC4B (nested within Subclade II-1) and VGC2B^824^ (Subclade II-2) [35,36]. The presence of *VdAve1* in a VC clade other than VCG2A (Subclade II-1) is in contrast to the recent finding of Jimenez-Diaz but consistent with earlier studies by de Jonge [26,27].

### UK strawberry isolates display a range of virulence on a common susceptible cultivar

In order to understand variation in virulence of *VdAve1*-containing isolates on strawberry, isolate pathogenicity was determined using a the susceptible cultivar of *Fragaria x ananassa.* Seven ‘race one’ *VdAve1-*containing isolates (two from UK hop, two from UK acer, three from Californian strawberry) and six ‘race two’ isolates (one from Californian strawberry and six from UK strawberry) were tested on the *V. dahliae* susceptible strawberry cv. Hapil. A wide range of symptoms were observed (Fig. 2a) ranging from full plant collapse to mild stunting. No clear association was observed between the presence of *VdAve1* and the virulence of the isolate by the end of the experiment, at six weeks post-inoculation (Fig. 2b).

**Fig 2.**
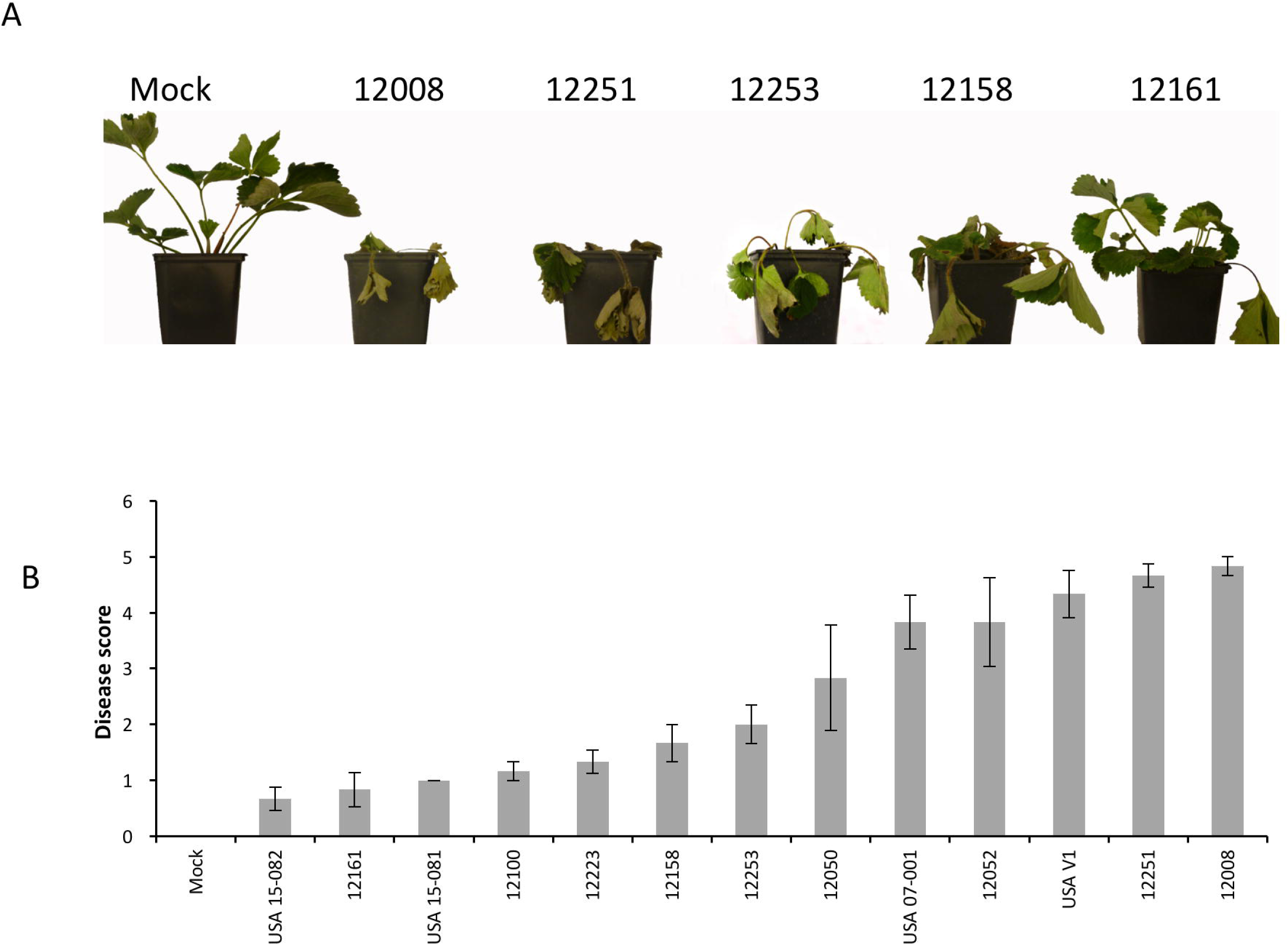
Indicative symptoms six weeks post inoculation of the strawberry cultivar ‘Hapil’ with *Vertcillium dahliae.* Disease scores six weeks post-inoculation (B) of the strawberry cultivar ‘Hapil’ inoculated with 13 isolates of *Verticillium dahliae.* Error bars are standard errors.

### VdAve1 isolates, when infected on strawberry have higher virulence early in the infection process and VC group is significantly associated with overall virulence level

A cumulative logit model was used to assess the severity of disease over time. From the results presented in Table 2, a clear pattern is apparent. *VdAve1* containing isolates across the VC groups display higher virulence early in the infection process, with the most pronounced effects at three weeks post inoculation. This variation is less visible as infection progresses and is undetectable at six weeks post-inoculation. Variation in pathogenicity associated with VC subclade grows in importance over time, with isolates in subclade II-2 displaying far higher pathogenicity, than isolates in subclade II-1. Isolate 12008 was found to be the most pathogenic isolate, with the highest level of disease recorded six weeks post inoculation (Fig. 2b). This isolate lacks *VdAve1* but is within subclade II-2.

**Table 2.** Likelihood ratio tests assessing effects of VC group and Race on pathogenicity

|  |  | Weeks post inoculation |  |  |  |  |  |  |  |  |  |  |  |
| --- | --- | --- | --- | --- | --- | --- | --- | --- | --- | --- | --- | --- | --- |
|  |  | 3 |  |  | 4 |  |  | 5 |  |  | 6 |  |  |
|  |  | Estimate of<br>antilogarithm<br>of fold<br>change | Std-Err | p Value | Estimate of<br>antilogarithm<br>of fold<br>change | Std-Err | p Value | Estimate of<br>antilogarithm<br>of fold<br>change | Std-Err | p Value | Estimate of<br>antilogarithm<br>of fold<br>change | Std-Err | p Value |
| Subclade II-2 |  | 1.6147 | 0.5980 | 0.00693 | 1.6290 | 0.5474 | 0.00292 | 2.1247 | 0.5692 | 0.000189 | 2.1082 | 0.5652 | 0.000192 |
| VdAve1 |  | 1.6924 | 0.6647 | 0.01090 | 1.1686 | 0.5344 | 0.02875 | 1.2533 | 0.5418 | 0.020713 | 1.0245 | 0.5461 | 0.060641 |

### The presence of VdAve1 alone is neither correlated with gain or loss of virulence on strawberry in artificial transformation of a highly virulent isolate

In order to examine whether the virulence of 12008 could be increased by the action of *VdAve1* we cloned the *VdAve1* gene from a ‘race 1’ isolate 12067 from tomato and then transformed the *VdAve1* expression vector into isolate 12008, a highly virulent isolate from UK strawberry using *Agrobacterium tumefaciens* mediated transformation. Vectors were constructed with either the inclusion of the native promoter region or use of a constitutive fungal promoter. In both cases five independent transformants (with the construct integrated at a random location in the genome) were selected for analysis. Strawberry plants inoculated with *VdAve1* knock-in conidia cultured in PDA media showed comparable wilt disease severity to the wild type (WT) isolate 12008 irrespective of whether the native promoter or constitutive promoter was used. Furthermore, even though qRT-PCR results confirmed that *VdAve1* knock-in mutants of PA4 with the native promoter and PG1 with a constitutive promoter had a similar *VdAve1* expression level with that of the ‘race 1’ isolate 12067, no significant difference was found between disease scores of the two mutants compared with the wild type isolate 12008 (Fig. 3a). qRT-PCR results with cultures of *VdAve1* knock-in mutants incubated on PDA or Czapek-Dox plates showed that *VdAve1* expression levels were induced (to differing degrees) on both PDA media and Czapek-Dox media (Fig. 3b). In conclusion the presence of *VdAve1* did not increase the virulence of *V. dahliae* on strawberry, when transformed into a virulent isolate.

**Fig 3.**
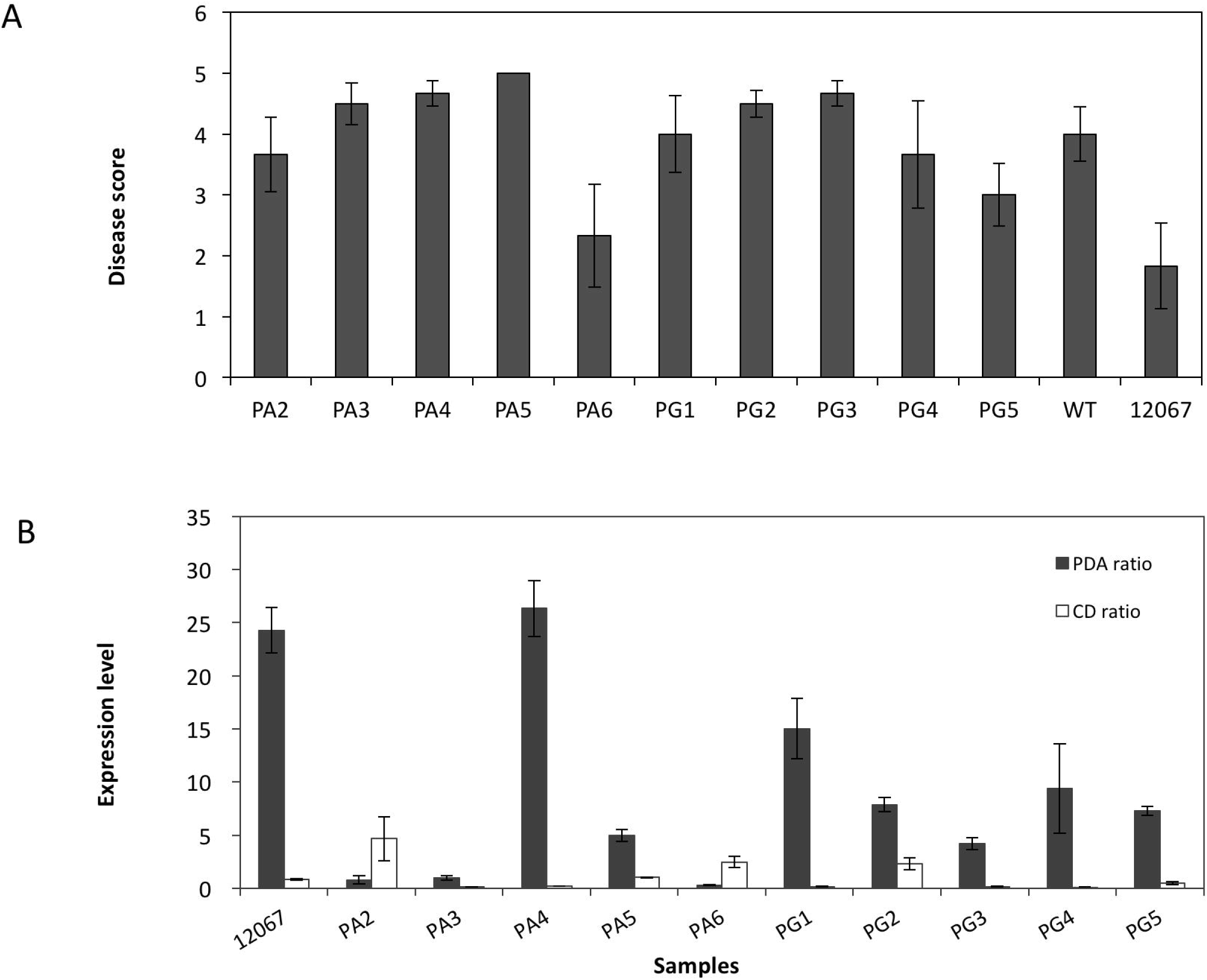
Disease symptoms of *Verticillium dahliae VdAve1* knock-in mutants generated in the wild-type isolate 12008 (A) on the susceptible strawberry cv. Hapil. Mutants labelled with ‘PA’ carry the *VdAve1* native promoter cloned from isolate 12067, whereas ‘PG’ mutants contained a constitutive promoter PgpdA from *Aspergillus nidulans*. The relative expression of *VdAve1* to the housekeeping genes elongation factor 1-α and beta tubulin in PDA and Czapek Dox agar cultures (B).

### Genome sequencing and assembly of selected isolates

To further characterise the molecular mechanisms of *V. dahliae* virulence, isolates from subclade II-1 and subclade II-2 were selected for genome sequencing. Long read PacBio and short read Illumina sequencing technologies were used to generate a reference genome for VC subclade II-1 isolate 12008. A further four isolates were sequenced using Illumina technology only, comprising two isolates from VC subclade II-2 (12158 and 12161) and two from VC subclade II-1 (12251, 12253).

High coverage was obtained for the isolate 12008 (174X), and 44-80x for the four Illumina-only isolates. A highly contiguous assembly was generated for isolate 12008 using 107-fold coverage of PacBio reads. *De novo* assembly using canu resulted in a 33.4 Mb assembly in 103 contigs (> 500 bp), with an N50 metric of 747 kb and a largest scaffold of 2.4 Mb (Table 3). RepeatModeler and RepeatMasker were used to identify repetitive and low-complexity regions within the 12008 assembly, masking 9.62 % (3.37 Mb) of the assembly. BUSCO was used to assess gene space, identifying that 3689 of 3725 (99 %) core Sordariomycete genes were present in the assembly. Gene prediction was carried out using Braker1 and CodingQuary, which predicted a total of 10440 genes in the unmasked assembly.

**Table 3.**
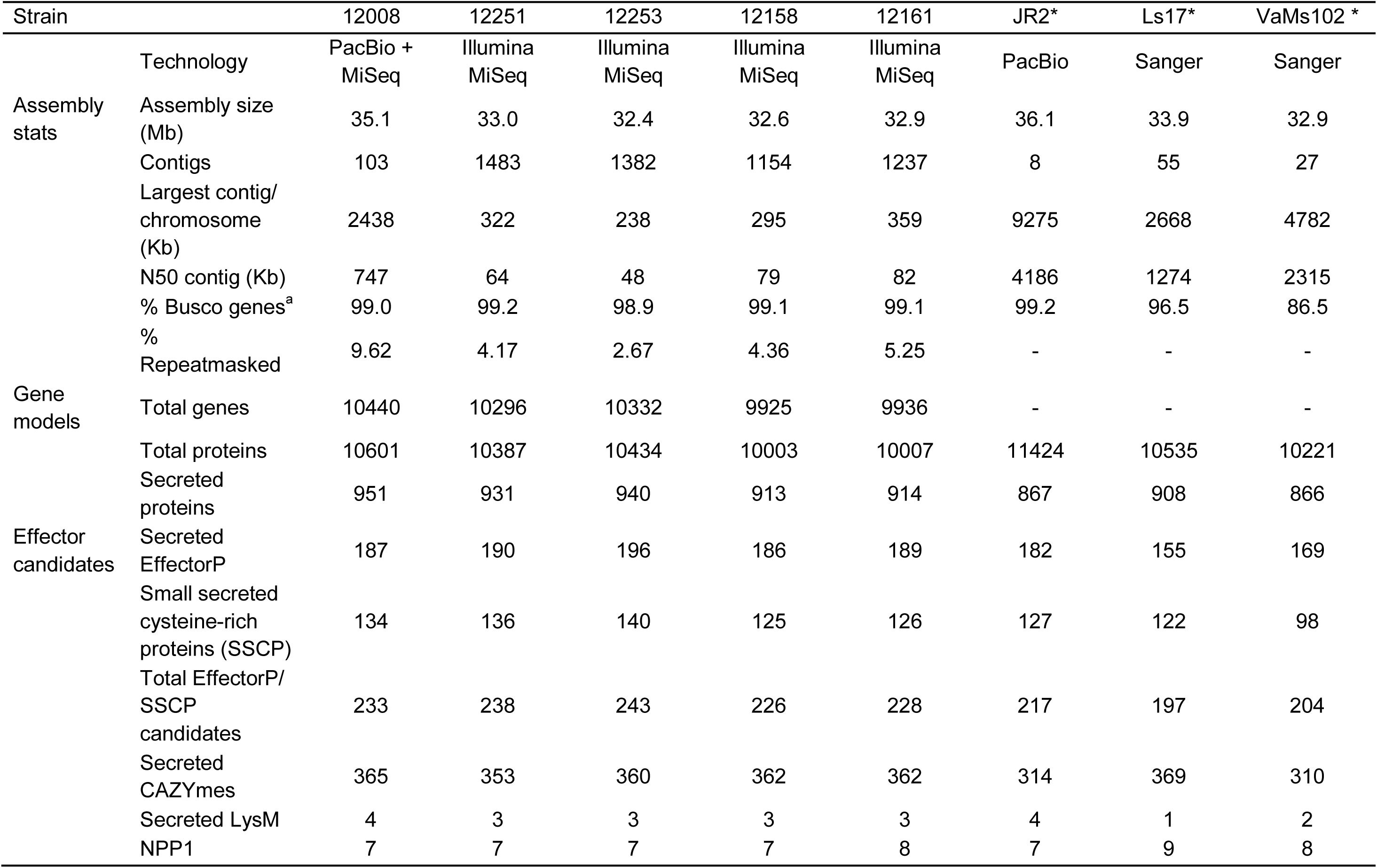
Genomic statistics of Verticillium strains.

*De novo* assembly of the four Illumina-only genomes resulted in assemblies of a similar size (32.4 - 32.9 Mb) and gene-space (98.9 - 99.2 % BUSCO genes) but more fragmented, with assemblies in 1154 - 1483 contigs (Table 3). Repetitive regions of the *V. dahliae* genome were not as complete in Illumina-only assemblies as the PacBio assembly, with 2.67 - 5.25 % of these assemblies masked. A total of 9925-10296 genes were predicted in these assemblies (Table 3).

Putative secreted effector proteins were identified using EffectorP and through searches for SSCPs. Results of effector prediction were broadly similar between the five sequenced isolates, with 187-196 EffectorP candidates and 125-140 SSCPs (Table 3). This led to a total set of 226-243 putative effectors predicted from each of the sequenced isolates. The number of secreted CAZymes in the five genomes ranged from 353-365. Seven chitin-binding lysin motif (LysM) proteins were identified in each proteome, apart from LS17 where eight were identified. Of the predicted LysM proteins, four were identified as secreted in the 12008 proteome and three predicted as secreted in the other sequenced isolates. Despite a similar total number of LysM proteins, reference proteomes carried different numbers of secreted LysM effectors (Table 3). Putative NPP1 (necrosis-inducing phytophthora protein) effectors were also identified, with isolate 12161 carrying an additional NPP1 protein that was not present in the other four isolates (Table 3).

### Vegetative compatibility groups have different effector complements and patterns of VdAve1 loss differ between VC groups

Even though the five isolates of 12008, 12251, 12253, 12158 and 12161 from UK strawberry were ‘race 2’ isolates, their virulence on strawberry was divergent as 12008 and 12251 can cause severe wilt disease while the isolate 12161 had very low virulence (Fig. 3A). After sequencing, short reads of each genome were mapped to the reference genome JR2 using bowtie, which is a *V. dahliae* ‘race 1’ isolate. The read coverage drop flanking *VdAve1* gene occurred earlier in the isolates of 12158 and 12161 than in isolates of 12008, 12251 and 12253 (Fig. 3B). Moreover, some reads from isolates 12158 and 12161 showed alignments at around 672 kb of the JR2 contig while no read coverage was observed over this region in the other three isolates (Fig. 4, panel A), indicating that these five ‘race 2’ isolates could be divided into two distinct groups.

**Fig 4.**
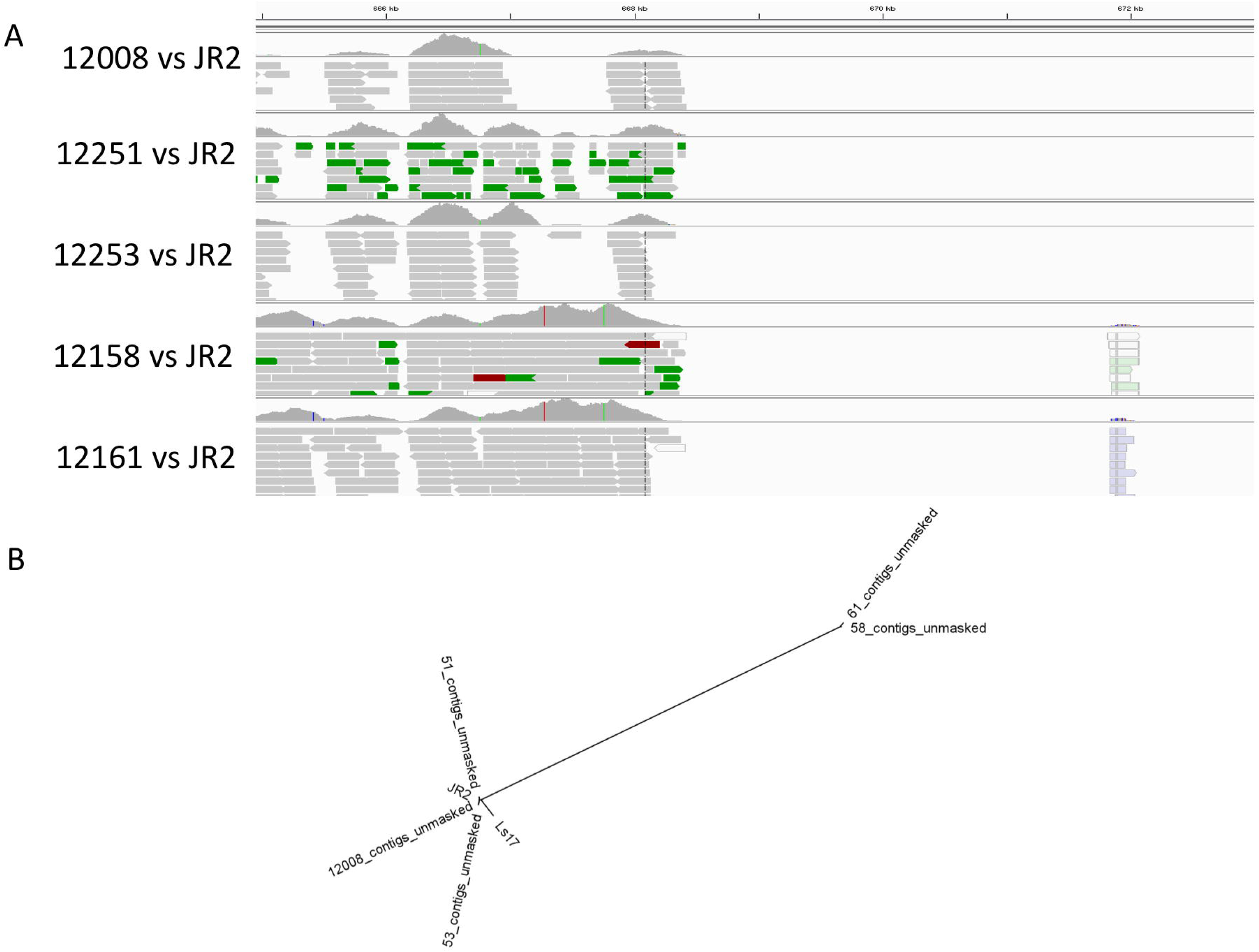
(A) Divergence of ‘race 2’ isolates from UK strawberry can be divided into 2 groups depending upon VC subclade. Alignment of short reads to the *VdAve1* region of the JR2 genome highlights different patterns of gene loss around the *VdAve1* region which are dependent upon VC subclade; Subclade II-2 isolates (12008,12251,12253) show similar patterns of read alignments to each other and subclade II-1 isolates (12158,12161) again show similarity within VC subclade but not between groups. (B) Whole-genome phylogenetic tree was generated using RealPhy based on the alignment of short or shotgun reads from each genome to the reference genome of JR2.

Phylogenetic analysis of RealPhy using whole-genome sequencing reads of these 5 ‘race 2’ isolates and the reference genome JR2 showed that *V. dahliae* isolates from UK strawberry should be divided into two clusters, consistent with the coverage mapping result. In the phylogenetic tree, the isolates of 12158 and 12161 were clustered together in a separate clade from the other isolates (Fig. 4, panel B). Interestingly, the isolates of 12008, 12251, 12253, JR2 and VdLs17 were clustered together (Fig. 4, panel B), which is concordant with the VC group analysis (Fig. 1).

### Effector expansion and losses in VC groups

To further identify those genes determining differences in virulence, orthology analysis was performed on the proteomes of the five UK isolates, *V. dahliae* reference isolates JR2 and VdLs17 and the *V. alfalfae* isolate VaMs102. In total, 80518 proteins clustered into 9697 orthogroups and a further 3110 proteins were considered orphan proteins, not clustering using OrthoMCL [60]. Results showed that the majority of proteins were conserved between *V. dahliae* and *V. alfalfae*, with 7285 orthogroups common to all isolates representing 81 % of the proteins (8542) from 12008. A further 501 orthogroups were conserved between all *V. dahliae* isolates, representing a further 5 % of proteins from 12008. A total of 361 orthogroups contained proteins predicted as effectors. Expansion of effector genes was investigated between VC subclades II-1 and II-2; 18 orthogroups were identified that showed patterns of gain or loss congruent with VC subclade (Table 4). Putative effector genes containing a glycoside hydrolase domain in orthogroup19 and a galactose oxidase domain in orthogroup73 were expanded in the higher virulence group isolates 12008, 12251 and 12253 (Table 2, Table 4), which may be responsible for aggressive performance. Nevertheless, effector genes containing the FAS1 domain in orthogroup700, necrosis inducing proteins in orthogroup736 and Zn (2)-C6 fungal-type DNA-binding domain in orthogroup824 were more abundant in the lower virulence isolates of 12158 and 12161 (Table 4, Supplementary Table 2).

**Table 4.** Variation in gene numbers of *Verticillium dahliae* strains in secreted and putative effector orthogroups

| Orthogroup | VC subclade II-2 |  |  |  |  | VC subclade II-1 |  | Annotation |
| --- | --- | --- | --- | --- | --- | --- | --- | --- |
|  | 12008 | 12251 | 12253 | Ls17<sup>a</sup> | JR2<sup>a</sup> | 12158 | 12161 |  |
| orthogroup19: | 10 | 9 | 9 | 10 | 9 | 7 | 7 | Glycoside hydrolase, catalytic domain |
| orthogroup73: | 7 | 7 | 7 | 4 | 5 | 1 | 1 | Galactose oxidase/kelch, beta-propeller |
| orthogroup104: | 4 | 4 | 3 | 3 | 3 | 3 | 3 | FMN-dependent alpha-hydroxy acid dehydrogenase domain |
| orthogroup509: | 4 | 2 | 3 | 1 | 4 | 0 | 0 | no annotation |
| orthogroup935: | 3 | 1 | 2 | 0 | 2 | 0 | 0 | no annotation |
| orthogroup1387: | 2 | 2 | 2 | 1 | 2 | 0 | 0 | no annotation |
| orthogroup631: | 2 | 2 | 2 | 1 | 2 | 1 | 1 | no annotation |
| orthogroup567: | 2 | 2 | 2 | 1 | 3 | 1 | 1 | no annotation |
| orthogroup8484: | 1 | 1 | 1 | 1 | 1 | 0 | 0 | Metallo-beta-lactamase |
| orthogroup8547: | 1 | 1 | 1 | 1 | 1 | 0 | 0 | Cyclin, C-terminal domain |
| orthogroup8589: | 1 | 1 | 1 | 1 | 1 | 0 | 0 | Transcription factor domain, fungi |
| orthogroup8712: | 1 | 1 | 1 | 0 | 1 | 0 | 0 | no annotation |
| orthogroup8757: | 1 | 1 | 1 | 0 | 1 | 0 | 0 | no annotation |
| orthogroup9016: | 1 | 1 | 1 | 0 | 0 | 0 | 0 | no annotation |
| orthogroup8586: | 1 | 0 | 0 | 1 | 3 | 0 | 0 | DPBB domain |
| orthogroup700: | 1 | 1 | 2 | 1 | 1 | 2 | 2 | FAS1 domain |
| orthogroup736: | 1 | 1 | 1 | 2 | 1 | 1 | 2 | Necrosis inducing protein |
| orthogroup824: | 1 | 1 | 1 | 1 | 1 | \|2 | 2 | Zn(2)-C6 fungal-type DNA-binding domain |

## Discussion

Our results clearly demonstrate that defining the clonal structure of Verticillium is key to understanding variability in pathogenicity and patterns of effector gain and loss. We conclude that VC subclade II-2 is associated with high virulence on the host plant strawberry, while VC subclade II-1, on the whole, contains isolates of lower virulence. To our knowledge this is the first time that variation in virulence on strawberry has been linked to VC group. However, comparable studies on the hosts artichoke and watermelon have also described an association of VC subclade II-2 with high virulence [61–63] indicating that this may be a common phenomena. High virulence was associated with VCG 1A on olive and cotton hosts, which may be explained by the presence of the high defoliating pathotype (D) within this lineage [64]; [65]. In contrast, no significant association was found between any VCG and virulence of *V. dahliae* on inoculated woody ornamentals, including Acer species [66]. We found no significant association between host of origin and variation in virulence on strawberry in our set of isolates (Table 2), which may suggest that the variation in virulence observed on some hosts (e.g. strawberry) that is associated with VC group may not be replicated in other hosts. This requires further study over a much larger number of isolates before a robust conclusion can be drawn.

Phylogenetic analysis of whole genome assemblies showed that the five sequenced isolates of *Verticillium dahliae* from UK strawberry clustered into two groups: the higher virulence isolates 12008, 12251 and 12253 and the lower virulence isolates, 12158 and 12161 in a separate clade adding support to the single locus separation seen when genotyping using the DB19/22 primers. This result suggests that the higher (VC subclade II-2) and lower virulence (VC subclade II-1) isolates are quite different across the whole genome. Even though these five isolates are defined as ‘race 2’ isolates, the group of isolates 12008, 12251 and 12253 were closely clustered with the ‘race 1’ isolate JR2, which indicated genetic diversity among ‘race 2’ isolates. It was recently reported that ten ‘race 2’ isolates can be divided into three subgroups depending upon the exact location of the read coverage drop when mapping the genomic reads to the ‘race 1’ isolate JR2 assembly [24]. Following this method, we found that the read depth of 12158 and 12161, surrounding the *VdAve1* gene dropped at the same positions when aligned to JR2, but this differed from the pattern of read dropout 12008, 12251 and 12253. It is therefore parsimonious to conclude that *VdAve1* was conserved before the split of *Vd* into VC subclade II-1 and VGC 2B and that there have been multiple independent loss events of *VdAve1* in these non-recombining, clonal lineages.

The plant cell wall provides the first barrier to prevent pathogen infection, and pathogens secrete numerous cell wall-degrading enzymes (CWDEs) to depolymerize the polysaccharide components of the plant cell wall [9]. Enzymes that are capable of hydrolytically cleaving glycosidic bonds in oligo- or polysaccharides (including cellulose and hemicellulose) are generally summarized under the term glycoside hydrolases [67,68]. Glycoside hydrolases are important plant cell wall degradation enzymes. In the current study, analysis of secreted and effector gene presence in orthogroups showed that genes containing domains of glycoside hydrolase and galactose oxidase were expanded in the high virulence isolate group of 12008 and 12251. Specifically, studies in the fungal genus Fusarium showed that plant galactose-containing hemicellulosic compounds may represent natural substrates of galactose oxidase [69]. The enriched domains of cell wall-degrading enzymes in isolates 12008, 12251 and 12253 may be responsible for greater degradation of plant cell wall composition and subsequent high virulence in *V. dahliae* isolates when infecting strawberry. This requires further testing to determine the causative elements responsible for the differences in virulence seen between the two VCG subclades. It also remains to be seen whether there is any difference in the resistance of cultivars to the two different clonal lineages of *V. dahliae*. Previous QTL mapping experiments in strawberry have been conducted under field conditions in highly infested plots which were artificially inoculated with many different culture of *V. dahliae [33]*. Further work using inoculations with single isolates is needed to understand whether resistance responses vary between isolates that differ in VCG.

A recent study showed that ‘race 2’ isolates are obtained with high frequency in many crops with the exception of several crops including strawberry in California, where ‘race 1’ isolates are found at a higher frequency [31]. This suggests that an ortholog of Ve1 is either absent or non-functional in the octoploid strawberry. However, when screening 27 *V. dahliae* isolates from UK strawberry, all of them were found to be non-*VdAve1*-containing ‘race 2’ isolates. Inoculation of common UK-grown wilt-susceptible cultivars of strawberry plants with ‘race 1’ isolates from other origins can cause wilt disease, indicating the lack of ‘race 1’ isolates on strawberry in the UK is not due to differences in host response between Californian and UK strawberry material. This indicates that Ve1-mediated resistance is not operating in the octoploid strawberry [20,70,71].

It is possible that geographical isolation plays a role in the frequency of *VdAve1* in the population. For example, all the artichoke isolates tested from California were ‘race 1’, though race 2 artichoke isolates from Tunisia have been reported, indicating that although the global migration of genotypes in *V. dahliae* has likely occurred and continues to play an important role in the diversification of fungal populations, within-field variation may be low and may be dominated by widespread clonal types. Thus, race structure in specific crops and commodities may exist, depending on the geography. Geography alone though does not fully explain the abundance of non-*VdAve1* containing isolates sampled from UK strawberry fields, as it is clear that *VdAve1*-containing isolates are present in the UK, as was revealed through extended sampling (Supplementary Fig. 1). It is of course possible that *VdAve1-* containing isolates have remained associated with their respective hosts and have not spread to infect strawberry growing regions of the UK. More detailed and systematic sampling would allow the spatial and host-associated distributions of *VdAve1-* containing isolates to be determined.

The results of pathogenicity assays using ‘race 1’ isolates from Californian strawberries revealed that *VdAve1* does enhance early development of pathogenic symptoms, the significance of which is attenuated as the disease progresses (Table 2). Although *VdAve1* was found to increase early symptom development in strawberry, whether present in VC subclade II-1 or VC subclade II-2, transformation of *VdAve1* either under its native promoter or under the *A. nidulans* promoter PgpdA failed to increase the virulence of the isolate 12008. Isolate 12008 is the most virulent isolate, based upon the pathogenicity test that we carried out (Fig. 2) and therefore we conclude that other factors present in the 12008 genome provide an alternative route to high virulence, such that the effect of *VdAve1* is masked.

Our work highlights that it is important that the nomenclature of *V. dahliae* is altered to move the classification of ‘race’ based on resistance responses in a single plant species, to one that is guided both by clonal lineage and effector complement as the adaptive potential of each lineage will differ, even if they contain the same effector.

## Acknowledgements

RF gratefully acknowledges the financial assistance of the China Scholarship Council (grant number: 201506300012). RJH was funded by BBSRC grants BB/K017071/1, BB/N006682/1, AA by BB/K017071/1, HC by innovate UK grant 101630. LA was funded by AHDB grant CP094. We would like to thank Vance Whitaker and Natalie Perez for Verticillium isolates. We also thank other members of the NIAB EMR (Andy Passey, Michelle Hulin, Beth Greenfield and Charlotte Nellist) for assistance throughout this project. We thank Bart Thomma and Luigi Faino for information about high-molecular weight DNA extraction.

RF and HC designed all experiments in collaboration with RJH. RF carried out all pathogenicity tests and transformations. HC carried out the VC group analysis. ECL assisted with genome assembly and annotation. AA provided support and supervision to RF for all informatics analysis, LA and HB generated Illumina genome sequences, XX provided statistical advice and assisted RJH with the development of the likelihood ratio tests. XH supervised RF and provided useful insights throughout the project. RF, HC, XX, AA and RJH wrote the manuscript. All authors approved the manuscript. The authors have no conflicts of interest to declare.

## Supplementary Figure Legends

**Supplementary Fig.1** PCR detection of *VdAve1* gene in *Verticillium dahliae* isolates from UK hop, acer, raspberry, potato, phlox, cotinus, and chrysanthemum (A) and strawberries (B) from UK and California. Primers target for V. dahliae intergenic spacer regions were used as control for DNA quality. The isolate of 12067 is a ‘race 1’ isolate as positive control and NTC was negative control of sterilized water.

